# Leaf cell wall properties and stomatal density influence oxygen isotope enrichment of leaf water

**DOI:** 10.1101/2020.10.15.341693

**Authors:** Patrick Ellsworth, Patricia Ellsworth, Rachel Mertz, Nuria Koteyeva, Asaph B. Cousins

## Abstract

Oxygen isotopic composition (Δ^18^O_LW_) of leaf water can help improve our understanding of how anatomy interacts with physiology to influence leaf water transport. Leaf water isotope models of Δ^18^O_LW_ such as the Péclet effect model have been developed to predict Δ^18^O_LW_, and it incorporates transpiration rate (*E*) and the mixing length between unenriched xylem water and enriched mesophyll water, which can occur in the mesophyll (*L*_m_) or veins (*L*_v_). Here we used two cell wall composition mutants grown under two light intensities and RH to evaluate the effect of cell wall composition on Δ^18^O_LW_. In maize (*Zea mays*), the compromised ultrastructure of the suberin lamellae in the bundle sheath of the ALIPHATIC SUBERIN FERULOYL TRANSFERASE mutant (*Zmasft*) reduced barriers to apoplastic water movement, resulting in higher *E* and *L*_v_ and, consequently, lower Δ^18^O_LW_. In cellulose synthase-like F6 (*Cslf6*) mutants and wildtype of rice (*Oryza sativa*), the difference in Δ^18^O_LW_ in plants grown under high and low growth light intensity co-varied with their differences in stomatal density. These results show that cell wall composition and stomatal density influence Δ^18^O_LW_ by altering the Péclet effect and that stable isotopes can facilitate the development of a physiologically and anatomically explicit water transport model.

## Introduction

The oxygen isotopic composition has been used to trace water transport into and through plants such as the identification of sources water taken up by a plant (Dawson & Ehleringer 1991; Ellsworth & Sternberg 2014) and water movement through the leaf (Barbour & Farquhar 2004; Barbour *et al*. 2017a). However, the use of oxygen isotopic composition to trace water transport is dependent on understanding the factors that influence water movement and the physical and biological processes that fractionate oxygen isotopes in water. For example, physiological factors such as transpiration (*E*) and leaf anatomical structures can influence water flow within the leaf and the degree of fractionation, impacting oxygen isotopic composition in leaf water [δ^18^O_LW_](Yakir 1998; Barbour & Farquhar 2000; Barbour *et al*. 2000a; Barbour 2007; Cernusak *et al*. 2016; Holloway-Phillips *et al*. 2016). Because of these complex interactions of δ^18^O_LW_-with leaf physiology and anatomy, it is presently difficult to draw clear conclusions from the experimental data (Barbour *et al*. 2017a). Therefore, a better understanding of how water movement within a leaf interacts with the physical and physiological processes must first be understood to be able to fully exploit δ^18^O_LW_(Cernusak *et al*. 2016; Barbour *et al*. 2017a).

Multiple models have been developed to help explain how environment and physiological processes influence δ^18^O_LW_. For example, the two-pool (Roden & Ehleringer 1999), the string-of-lakes models (Gat & Bowser 1991), and the Péclet effect (Barbour*et al*. 2000b; Barbour *et al*. 2004) have been developed to help explain how evaporatively-enriched water at the sites of evaporation and unenriched δ^18^O_sw_ influence δ^18^O_LW_. In all of these models, unenriched water supplied through the xylem becomes enriched at the sites of evaporation based on a modified Craig-Gordon model, where enrichment is a function of relative humidity, leaf temperature, and oxygen isotopic composition of vapor (See Theory section; Craig & Gordon 1965; Farquhar *et al*. 1998). The two-pool model operates under the principles of mass balance where unenriched water in the xylem and enriched water from the sites of evaporation (δ^18^O_e_) are considered two discrete water pools that mix to form δ^18^O_LW_. Alternatively, the string-of-lakes model used in grasses considers that instead of two discrete water pools there are multiple discrete pools in series up the leaf blade, where the source water to each pool is the water from the pool preceding it. This leads to a progressive enrichment of water from the first pool at the base of the leaf blade to the last pool towards the leaf tip. However, the two-pool and the string-of-lakes models do not address cases when δ^18^O_LW_ varies with *E* (Farquhar & Lloyd 1993).

The Péclet effect model accounts for the continuous gradation from unenriched water moving through the xylem to enriched water at the sites of evaporation through opposing flows of advection and back-diffusion (Barbour *et al*. 2000a; Barbour *et al*. 2000b). These opposing flows in the Péclet model are dependent on *E* and the effective mixing length over which unenriched water from the xylem mixes with enriched water from the sites of evaporation (*L*). Lcan be unique to each tissue type where these two isotopically-different water pools are mixing, such as in the mesophyll as water moves from the xylem to the inner-stomatal cavity [*L*_m_] or in the veins where enriched mesophyll water mixes with unenriched xylem water [*L*_v_] (Holloway-Phillips *et al*. 2016). In this manner, isotopic mixing occurs radially from the xylem out to the stomata and longitudinally in the veins where enriched water enters the veins through back diffusion and is carried toward the leaf tip (Farquhar & Gan 2003). Leaf traits such as cell wall composition that can affect the extent that mixing occurs by influencing the length and tortuosity od the path that water takes through the xylem and mesophyll (*L*) and flow or conductivity (*E*), which, in turn, affect δ^18^O_LW_ beyond the effect that δ^18^O_SW_ has. Each of these models have found empirical support under specific conditions and for certain species; however, none of them can universally explained the isotopic behavior of leaf water (Roden & Ehleringer 1999; Barbour & Farquhar 2000; Barbour *et al*. 2000b; Helliker & Ehleringer 2000; Barbour & Farquhar 2004; Loucos *et al*. 2015; Song *et al*. 2015; Holloway-Phillips *et al*. 2016).

However, the Péclet effect model is unique in that it can be used to test hypotheses on how leaf anatomical features affect the movement of leaf water and influence δ^18^O_LW_. Specifically, this model can be used to test how leaf traits influence the advection of water from the xylem to the sites of evaporation and back diffusion of water from the sites of evaporation into the mesophyll and xylem, which determines the influence of the enrichment of water at the sites of evaporation above source water (Δ^18^O_e_) on leaf water enrichment above source water (Δ^18^O_LW_). In other words, anatomical features that potentially influence the mixing of water pools and the flow of water out of the leaf (*E*) or that impact Lcan be tested on how they alter Δ^18^O_LW_. For example, greater vein density increased the oxygen isotopic enrichment along the grass leaf blade because it increased mixing of enriched mesophyll water with xylem water, leading to the progressive enrichment of xylem water and Δ^18^O_e_ up the leaf blade and subsequently Δ^18^O_LW_(Helliker & Ehleringer 2000; Farquhar & Gan 2003). In dicots, greater vein density had the opposite effect in that Δ^18^O_LW_ was lower because the proportion of unenriched xylem water (*f*_sw_) was greater (Holloway-Phillips *et al*. 2016). Stomatal density and pore size also influence Δ^18^O_LW_, where greater stomatal densities, even when E remained the same, had higher Δ^18^O_LW_ because L decreased with stomatal density (Sternberg & Manganiello 2014; Larcher *et al*. 2015; Liang *et al*. 2018). These findings have led to increased interest in anatomical features such as cell wall composition that partly control the permeability of the cell wall and, in turn, influence the relative importance of apoplastic water movement from the xylem to the sites of evaporation, which ultimately affectsE and L (Barbour *et al*. 2017b).

Cell wall composition mutations modify how leaf anatomy influences water movement andphysiological properties that affect *E, L*, and, ultimately, Δ^18^O_LW_. For example, suberin in the suberin lamellae provides a barrier to apoplastic water movement and passive water loss from the vasculature (Mertz & Brutnell 2014; Vishwanath *et al*. 2015). *Zmasft* double mutants, which were created in maize (*Zea mays* L.) by mutating two paralogously duplicated, unlinked maize orthologues of Arabidopsis thaliana *ALIPHATIC SUBERIN FERULOYL TRANSFERASE*, have a compromised suberin llamelae ultrastructure that increased cell wall elasticity and apoplastic water diffusion (Mertz *et al*. In review). Greater apoplastic diffusion and cell wall elasticity can increase available water for transpiration, while less barriers to water movement can increase the number of water channels and modify the relative importance of each channel that water takes in and out of the leaf vasculature and from the vasculature to the stomata, influencing *L*_m_ and *L*_v_. Cellulose synthase-like F6 (*CslF6*) in rice (*Oryza sativa* L.) mediates the biosynthesis of mixed linkage glucan (MLG), a polysaccharide important in cell wall composition (Vega-Saénchez *et al*. 2012). Loss-of-function mutants do not synthesize MLG (Vega-Sénchez *et al*. 2012; Smith-Moritz *et al*. 2015) and have higher cell wall porosity, which has been show to affect the internal conductance of CO_2_ (*g*_m_) (Ellsworth *et al*. 2018). This increase in porosity affected the movement/conductance of CO_2_ and may also alter apoplastic water movement as well, which can alter water flow to the stomates (*E*) or the length and tortuosity of water path (*L*). Therefore, these mutants provide a means to evaluate the effect of cell wall composition on leaf physiology with respect to water transport through its impact on Δ^18^O_LW_.

To determine the influence of cell wall properties on Δ^18^O_LW_ we tested how the loss of MLG and suberin changed water movement from the xylem to the sites of evaporation under environmental conditions (growth light and relative humidity) that have been demonstrated to influence how *E* and *L* impact Δ^18^O_LW_. The data presented here demonstrates that changes in both cell wall properties and stomatal density influence Δ^18^O_LW_ through the impact of *E* and *L* on the Peclet effect.

### Theory and calculation of isotopic factors

Leaf water isotopic composition (δ^18^O_LW_) is a mixture of two water pools that represent the sources of isotopic variation within the leaf source water in the xylem (δ^18^O_SW_) and water from the sites of evaporation (δ^18^O_e_ Equation 1), where using a mass

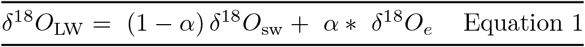

balance equation *α* is the fraction of unenriched water in leaf xylem. To remove the effect of source water on δ^18^O_LW_, the oxygen isotopic composition in leaf water can be expressed as enrichment above source water (Δ^18^O_LW_),

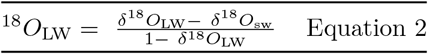

where δ^18^O_LW_ and δ^18^O_sw_ are expressed as ratios of ^18^O/^16^O (Barbour 2007; Cernusak *et al*. 2016). Therefore, eqn. 1 can be expressed as enrichments (Eqn. 3).

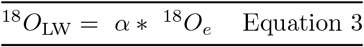

Water at the sites of evaporation is assumed to follow the Craig-Gordon model, so Δ^18^O_e_ can be derived from equilibrium fractionation (ε^+^), the ambient to intercellular air vapor mole fraction (*e*_a_/*e*_i_), the kinetic fractionation (ε_k_), and air vapor enrichment (Δ^18^O_v_; Eqn. 4) (Craig & Gordon 1965; Dongmann *et al*. 1974; Flanagan *et al*. 1991).

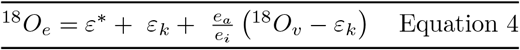

ε^+^ is the fractionation associated with the movement of water from the liquid phase into the vapor phase as part of the boundary layer around the leaf surface and is inversely related to T_L_ (Eqn. 5).

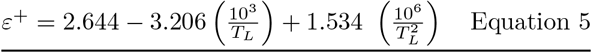

The fractionation (*ϵ*_k_) associated with water vapor leaving the boundary layer is inversely proportional to the stomatal (*g*_s_) and boundary layer (*g*_bl_) conductances (Eqn. 6).

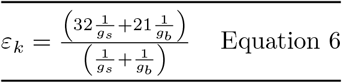

Δ^18^O_LW_ is nearly always less enriched than Δ^18^O_e_, meaning that leaf water is not a well-mixed pool undergoing evaporative enrichment, so two models were developed to explain Δ^18^O_LW_ based on its isotopic sources: the two-pool model (Eqn. 1) and the Peclet effect model (Roden & Ehleringer 1999; Barbour & Farquhar 2000; Helliker & Ehleringer 2000; Gan *et al*. 2003; Barbour 2007). The Péclet effect model, instead of the fraction of water contributed by the sites of evaporation (α in Eqn. 3) being fixed, it was modified to be dependent on the Péclet number (P; Eqn. 7). The Péclet number

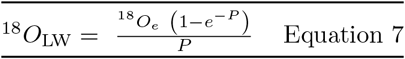

represents the advective water flow from the xylem toward the sites of evaporation and back diffusion from the sites of evaporation into the rest of the leaf including the xylem.

The Peclet number is proportional to the transpiration rate (*E*) and the effective mixing length (*L*) that water takes to move from the veins to the sites of evaporation, and inversely proportional to the molar density of water (*C*; 5.56 x 10^-4^ mol m^-3^) and the diffusivity of H_2_^18^O in water (*D*; Eqn. 8) (Farquhar *et al*. 1998; Barbour & Farquhar 2000; Barbour *et al*. 2000b; Gan *et al*. 2003; Barbour & Farquhar 2004; Barbour 2007). The diffusivity of H_2_^18^O in water calculated according to Cuntz *et al*. (2007).

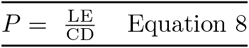

*L* is the product of two components related to leaf anatomy: one that accounts for the tortuosity of water pathway from the xylem to the stomatal pore (*l*) and the other refers to the cross-sectional area perpendicular to water flow relative to the leaf surface area (*k*) (Barbour & Farquhar 2004; Cernusak & Kahmen 2013; Song *et al*. 2013; Sternberg & Manganiello 2014). The assumption that *L* is constant for species, across treatments, and across a gradient of transpiration rates has not always been supported (Song *et al*. 2013; Roden *et al*. 2015). The lack of understanding of *L* and the factors that affect it largely arise from the difficulty of measuring *L*.

The determining evidence of the presence of the Péclet effect is if the fractional contribution of source water (unenriched xylem water; *f*_sw_) increases with *E* (Eqn. 9; Song *et al*. 2015; Holloway-Phillips *et al*. 2016). A Péclet effect has failed to be observed in several

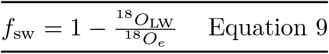

studies and species (Roden & Ehleringer 1999; Cernusak *et al*. 2003; Loucos *et al*. 2015; Song *et al*. 2015; Holloway-Phillips *et al*. 2016). Adherence to the Péclet effect model appears to depend on the relative importance of enriched vein water to bulk leaf water and the interaction between xylem and mesophyll water pools (Song *et al*. 2015; Holloway-Phillips *et al*. 2016).

Leaf anatomy was shown to play a distinct role in Δ^18^O_LW_, where parallel venation in grasses increased the magnitude of enrichment from the base of a grass leaf blade to the tip (Helliker & Ehleringer, 2000). Even though Δ^18^O_e_ remains the same over the length of the leaf blade, δ^18^O_SW_ (xylem water) becomes progressively more enriched up the blade, supplying increasingly more enriched water to the sites of evaporation and creating a gradient in δ^18^O_LW_. Vein density increased the influence of the enriched mesophyll water on the xylem, increasing the magnitude of δ^18^O_LW_ along the blade. The string-of-lakes model described this progressive enrichment up the leaf blade as a series of interconnected pools where the source of one pool is the preceding pool (Gat & Bowser 1991; Helliker & Ehleringer 2000). The Farquhar-Gan model was created, so that separate Péclet effects were associated with xylem and mesophyll (lamina) water pools (Eqn. 10; Gan *et al*. 2003; Holloway-Phillips *et al*. 2016).

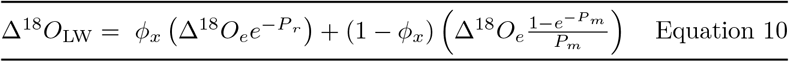

Where x is the fraction of leaf water in the xylem, and P_r_ and P_m_ are the xylem and mesophyll Peclet numbers, respectively. This model allows for different Peclet effects in each tissue type (Gan *et al*. 2003; Holloway-Phillips *et al*. 2016). The influence of the xylem Péclet depends on physiological traits such as *E* that regulates flow velocity and anatomical traits, such as vein density, fraction vein volume, stomatal density and size, and specific water content (Sternberg & Manganiello 2014; Larcher *et al*. 2015; Song *et al*. 2015; Holloway-Phillips *et al*. 2016; Liang *et al*. 2018). Potentially multiple other anatomical traits that influence the movement of water along the transpiration pathway and in the leaf alter Δ^18^O_LW_. The Farquhar-Gan model can be modified by excluding the mesophyll Péclet if it is small or nonexistent compared to xylem Péclet (Eqn. 11). Therefore, the Péclet effect

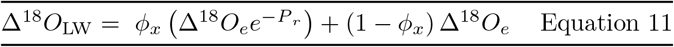

model can be adapted to the species depending on the leaf anatomy and physiology influencing Δ^18^O_LW_.

## Materials and Methods

### Plant Material

#### Experiment 1: Maize suberin mutants (Zmasft)

Maize (*Zea mays* L.) seeds were planted in 11.3 L pots and placed in a controlled-environment growth chamber (Biochambers, GRC-36, Winnipeg, MB, Canada). The photoperiod was 16 h with a 1-hour ramp for light reaching 1000 μmol quanta m’V at canopy level and 1-hour ramp for temperature to mimic dawn and dusk. Day/night temperatures were set at 28°C/20°C, respectively and relative humidity was kept at 70 %. The pots were distributed in two light treatments even before germination: high light (HL) exposed to 1200 μmol quanta m^-2^s^-1^ at canopy level and low light (LL) exposed to 300 μmol quanta m^-2^ s^-1^ at canopy level. The plants were grown simultaneously in the same growth chamber and the height of the pots was adjusted to maintain the desired light treatments throughout the experiment.

Five replicates of each genotype were grown in a soil mixture of Sunshine Mix LC-1 (Sun Gro Horticulture). Plants were watered daily and fertilized every four days, initially with 20-20-20 and later with 15-5-15 CalMag. Experiment was conducted from March to May 2015, at Washington State University, Pullman, WA, USA.

#### Experiment 2: Rice mixed linkage glucan mutants (cslf6)

Rice (*Oryza sativa* L.), Nipponbare cv (wild type) and mutants*cslf6-1* and *cslf6-2*, which vary in the position of their transposon insertion within the *CslF6* genomic sequence of Nipponbare cv(Vega-Sénchez *et al*. 2012; Ellsworth *et al*. 2018) were germinated on wet filter paper in a petri dish. Seedlings were transplanted in trays containing Sunshine Mix LC-1 soil (Sun Gro Horticulture) and turface (ratio of 3:1 *v*/*v*) and then after a month, they were transplanted into 7.5 L pots and placed in a controlled environment growth chamber (Biochambers, GRC-36, Winnipeg, MB, Canada). The photoperiod was 14 h with a two-hour ramp, reaching 500 μmol photon m^-2^ s^-1^ at canopy level. Day/night temperatures were set at 26/22 °C, respectively with a one-hour ramp, and relative humidity was kept at 70 %. One week after transplanting, plants were distributed into two light treatments: high light (HL) plants were submitted to a gradual increase in irradiance over 5 days to 1000 μmol photon m^-2^ s^-1^ at canopy level and low light (LL) plants were placed directly under 300 μmol photon m^-2^ s^-1^ at canopy level. Plants were grown simultaneously in the same growth chamber, and pot heights were adjusted to maintain the desired light treatments at canopy level.

Five replicates of each genotype were watered daily and fertilized every four days with a nutrient solution containing Sprint 330 iron chelate (1.86 g L^-1^), magnesium sulfate (0.88 g L^-1^), Scotts-Peters Professional 10-30-20 compound (3.96 g L^-1^), and Scott-Peters Soluble Trace Element Mix (10.0 mg L^-1^; Scotts). The experiment was conducted from March to May 2014, at Washington State University, Pullman, WA, USA.

### Anatomical analysis

For measurements of partial volume of total vasculature, vascular bundles (both including and excluding the bundle sheath), and xylem, cross sections were made on leaf 5 at the point of maximum lamina width. The freehand transverse sections were immersed in water and imaged under UV light filter (with excitation wavelength range 350-520nm) using cell wall autofluorescence to maximize the contrast between the bundle sheath cell wall and the surrounding mesophyll cells. Images were captured using Leitz Epi-Fluorscent Microscope with Leica DFC425C Camera at 100x for the leaf blade and 250x for the midrib. Additional images were taken at 40x magnification for the leaf blade and 25x for the midrib to calculate cross sectional leaf area. The section area of all vascular bundles (including and excluding the bundle sheath) was measured. For xylem area, diameter of each xylem element was measured twice perpendicular to each other, and mean diameter was used to calculate xylem area as the area of a circle. All measurements were made in ImageJ (Schneider 2012). The total vasculature was calculated as area of the vascular bundles in the leaf blade plus the entire midrib. Partial volume of each was calculated as the total cross-sectional area of each respective anatomical feature relative to the total leaf cross-sectional area (blade area + midrib area).

The surface of three fresh leaves per genotype per treatment was analyzed under the low vacuum mode of FEI Scanning Electron Microscope (SEM) Quanta 200F (FEI Company, Field Emission Instruments, Hillsboro, OR, USA). The leaves were the same or similar to those used for gas exchange measurements and anatomical analyses. Images of the adaxial and abaxial epidermal surfaces were randomly captured across the entire leaf blade avoiding the central vein (min. 10 images per leaf) to determine the number of stomata per mm^2^ of leaf surface (stomatal density) and axis length of stomatal pores.

### Gas Exchange Measurements

Gas exchange measurements were performed in Pullman, WA, USA, with mean atmospheric pressure of 92.1 kPa. The mid portion of 3-5 uppermost, fully expanded leaves were used for gas exchange measurements using a LI-6400XT and a 2 x 3 cm leaf chamber with a red and blue light emitting diode light source (P/N 6400-02B, Li-Cor, Lincoln, NE, USA). Spot measurements were performed inside the growth chamber, and the light intensity, temperature, and relative humidity during gas exchange measurements were maintained similar to the growth chamber conditions under each treatment. Leaf-to-air vapor pressure deficit was kept at 1 kPa, flow rate maintained at 300 μmol air s^-1^, and oxygen partial pressure of 19.3 kPa (21 %).

### Leaf trait measurements

The fully expanded leaf selected for transpiration measurements (section above) was afterwards collected, photographed for calculation of area using ImageJ (Schneider 2012) and placed in weighed glass tubes. Fresh weight was first recorded and its water extracted according to Vendramini and Sternberg Lda (2007) for measurements of δ^18^O_LW_. The dried leaf material was weighed to calculate leaf water content (LWC) and leaf dry mass per area (LMA). LWC was calculated as the difference between fresh and dry leaf weight per area and LMA was calculated as dry leaf weight per leaf area.

Root crowns were collected and placed in a separate glass tube for water extraction (Vendramini & Sternberg Lda 2007) and oxygen isotope analysis, so that source water was characterized for each plant to calculate leaf water oxygen isotope enrichment (Δ^18^O_LW_). Vapor from the growth chamber was collected every 30 min during transpiration measurements. The vapor was sealed in 5 L inert foil gas sampling bags (Supelco Analytical #30228-U; Bellefonte, PA) that restrict exchange of water vapor or CO_2_ exchange with ambient air, and after collection the vapor samples were measured immediately (details below).

### Oxygen isotope analysis

Root crown and leaf water oxygen isotope ratios (δ^18^O_LW_) were analyzed by isotope ratio mass spectrometer (IRMS). A 0.3% CO_2_ /He gas mixture was flushed through the vials and equilibrated with water for 48 h on a ThermoFinnigan GasBench II (Bremen). The equilibrated CO_2_ gas was separated at 40 °C with a 25 m x 0.32 mm ID GC column (Varian, poraplot Q) and analyzed with a continuous flow isotope ratio mass spectrometer (Delta PlusXP, Thermofinnigan, Bremen) (Brenna *et al*. 1997; Qi *et al*. 2003) to derive the oxygen isotope ratios of the water. Standards (SLAP and Puerto Rico water) were analyzed alongside the samples. Final delta values were the mean of 5 sample peaks reported relative to Vienna Standard Mean Ocean Water (V-SMOW).

Irrigation water and water vapor were measured by isotope ratio-cavity ringdown spectroscopy using a water analyzer (model L1102-i, Picarro, Sunnydale, CA, USA). Each sample was analyzed six consecutive times, and the mean of the last three analyses was used. Three laboratory standards, calibrated to the V-SMOW scale, were interspersed among the samples and used to correct the sample δ^18^O values to the V-SMOW scale. Water vapor was introduced into the water analyzer through the inlet valve and analyzed for at least 15 minutes, and the δ^18^O of water vapor (δ^18^O_v_) was calculated as the mean δ^18^O over the last five minutes. The δ^18^O_v_ was corrected to the V-SMOW scale in the same way that irrigated water samples were corrected. Further δ^18^Ov was corrected for water concentration dependency. The stable isotope composition of oxygen (δ^18^O) was reported in δ notation (Eqn. 12),

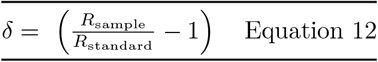

where R_sample_ and R_standard_ are the isotopic ratios of oxygen (^18^O/^16^O) of the sample and international standard, respectively. The precision of the analysis for each parameter was 0.2 and water vapor and 0.07

For calculation of effective leaf water mixing length, some parameters were measured, and others were calculated. Steady state condition (SS) was assumed for all calculations. Water isotope enrichment at the sites of evaporation (Δ_e_) was calculated for each leaf (Craig & Gordon 1965; Dongmann *et al*. 1974) relative to source water (δ^18^O_e_ - δ^18^O_sw_; Eqn 2). Δ^18^O_v_ is the atmospheric water vapor enrichment above plant water source (δ^18^O_v_ - δ^18^O_sw_), which in this study was root crown water. We used Δ^18^O_v_ from the atmosphere, as only 6.5 % of the leaf was inside the cuvette during gas exchange measurements for about 3-5 min.*e*_a_/*e*_i_ is the ratio of vapor pressure external to the leaf to internal vapor pressure, ε* is the temperature-dependent (°Kelvin) equilibrium fractionation factor between vapor and water inside the leaf (Eqn. 5; Majoube 1971), and ε_k_ is the kinetic fractionation during diffusion of vapor from the leaf to the atmosphere (Eqn. 6; Farquhar *et al*. 1989).

### Statistical Analysis

All statistical analyses and graphing were performed in R software (version 3.2.2) (R Core Team, 2015). The number of replicates varied depending on the measured parameters. Leaf anatomy study was performed in 3-5 replicates. The remaining measurements were performed in 5 replicates. If the dependent factor was not normal or homogeneous, the data were transformed before using a two-way ANOVA. Two-way ANOVA followed by contrasts was selected to estimate mean differences (LL plants subtracted from HL plants) for each genotype separately for individual parameters. Genotypes were combined for the ANCOVA followed by contrast analysis to determine if changes in leaf water oxygen isotope ratios co-varied with changes in other physiological and anatomical traits (in terms of mean differences of HL and LL growth treatment). Contrast analysis was considered the appropriate statistical approach to interpret mean differences according to personal consultation provided by the statistical facility at WSU.

According to Péclet effect theory, the extent to that the Péclet effect is occurring varies with leaf water pool, so the relative importance of each Péclet effect is dependent on its strength and water pool size (Farquhar & Gan 2003; Holloway-Phillips *et al*. 2016). The two Péclet models fitted to the *f*_sw_ versus *E* data were the xylem and the mesophyll and xylem Péclet models where the unknowns were *L*_v_ and x in the xylem Péclet model and *L*_m_, *L*_v_, and x for the mesophyll and xylem Peclet model. The knowns were *f*_sw_ (1 - Δ^18^O_LW_/Δ^18^O_e_) and *E*. Model fitting was done with the Levenberg-Marquardt Nonlinear Least-Squares Algorithm in R using the minpack.lm package.

## Results

### Maize Zmasft double mutant versus wildtype

The enrichment of water at the sites of evaporation above source water (Δ^18^O_e_) was 4.1 times more responsive to changes in RH than to changes in light intensity, meaning that *e*_a_/*e*_i_ is a more influential variable in determining Δ^18^O_e_ than *g*_s_-driven changes in ε_k_. Δ^18^O_e_ did not differ between genotypes in either RH or light intensity, while maize *Zmasft* double mutants were less enriched in leaf water oxygen isotope enrichment (Δ^18^O_LW_) than wildtype in every treatment (14.24 ± 1.48 and 16.14 ± 1.41 1a, b). In all treatments, Δ^18^O_LW_ was consistently less enriched in the *Zmasft* double mutants than wildtype by 0.5 to 1 Δ^18^O_e_ values trended lower in wildtype, contrary to what is expected if differences in Δ^18^O_e_ were driving the genotypic differences in Δ^18^O_LW_. Nonetheless, Δ^18^O_e_ has a drastic effect on Δ^18^O_LW_ in that Δ^18^O_LW_ was 11.65 ± 0.65 % RH than at 50 % RH, ranging from 9.44 to 22.80 light intensity was much less in that Δ^18^O_LW_ was 2.90 ± 0.50 than at 1200 μmol photons m^-2^ s^-1^, and there was no difference at 80 % RH (Table 1, S1; Fig. 1).

**Fig. 1.**
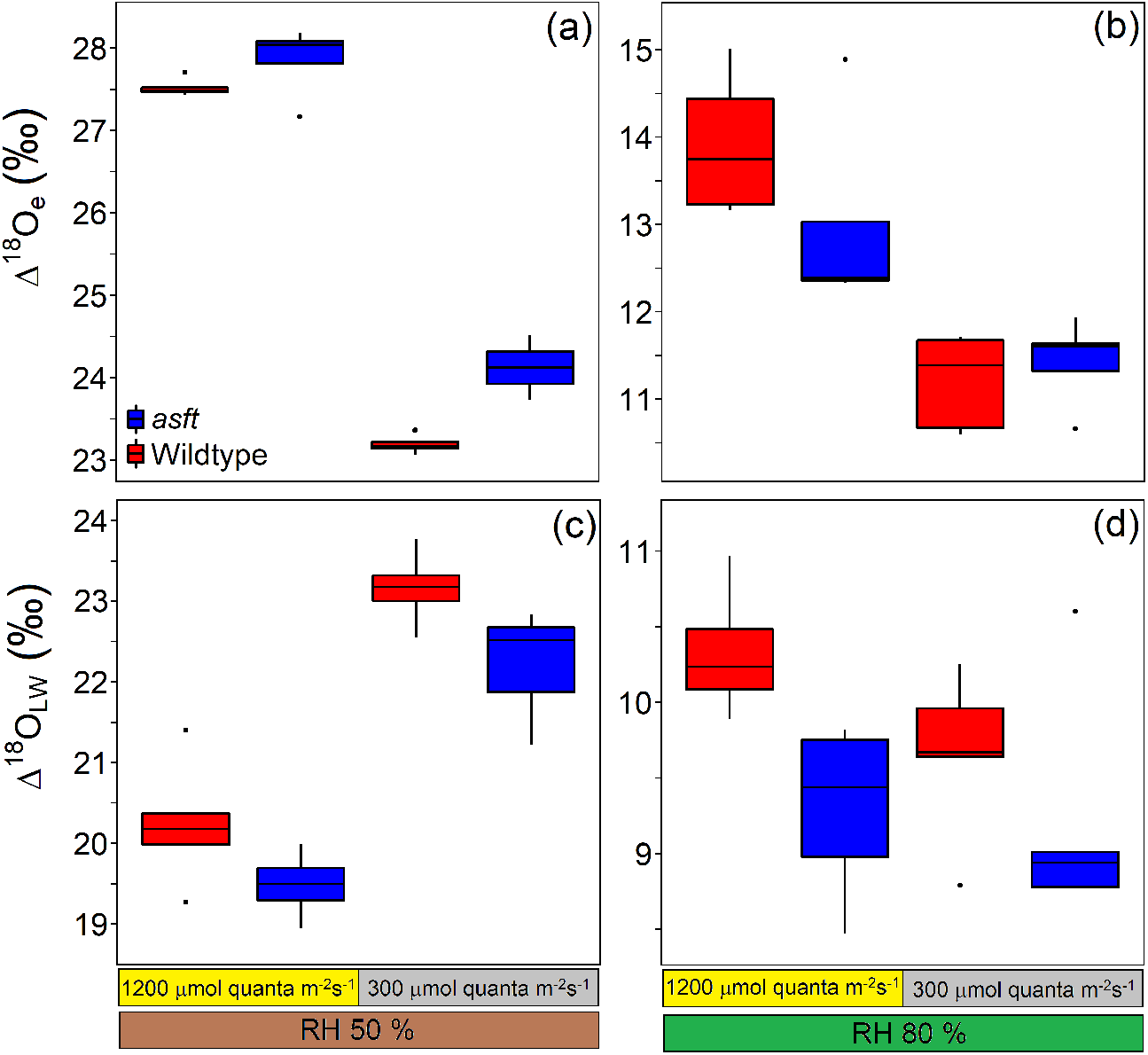

**Table 1.**
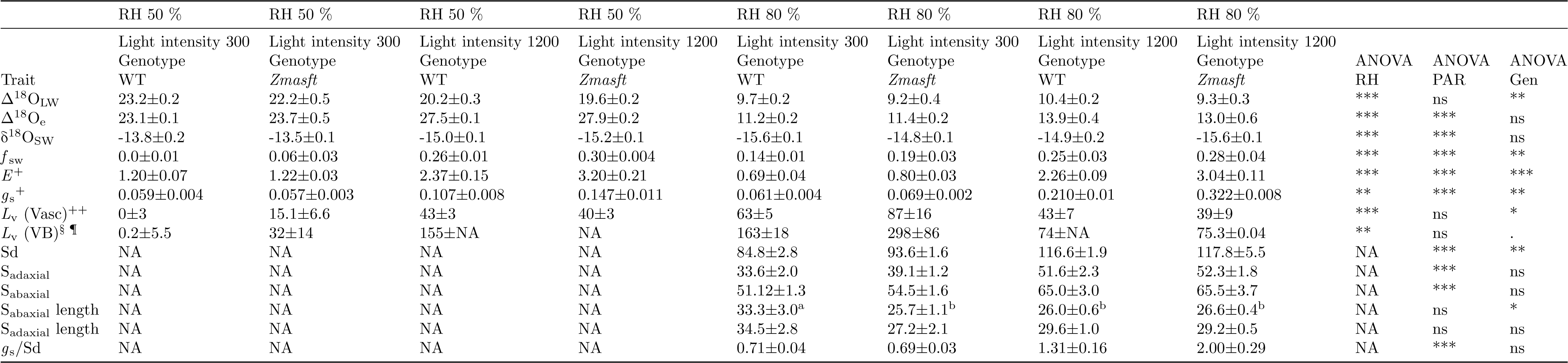
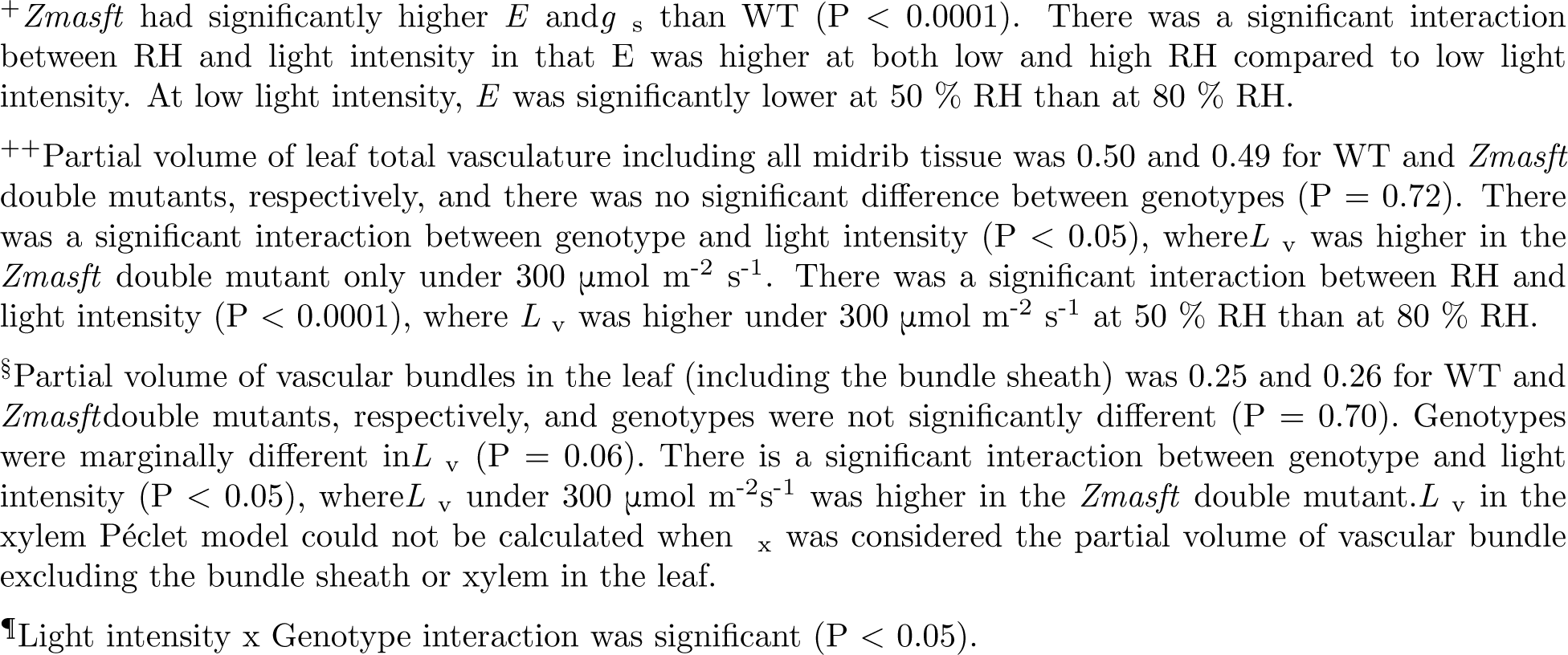
Leaf water isotope parameters for *Zea mays* wildtype and Zm*asft* double mutants (aliphatic suberin feruloyl transferase), grown under 300 or 1200 photosynthetic active photon flux density (PAR; μmol photons m^-2^ s^-1^) and either 50 or 80 % relative humidity (RH). Stomatal measurements were on plants grown at 80 % RH. Results of the main effects of a three-way ANOVA are included, except the stomatal traits. Stomatal traits were analyzed using a two-way ANOVA because they were only measured in plants grown under 80 % RH. Significance was shown as ns, *, **, and *** for not significant, P < 0.05, P < 0.01, and P < 0.001, respectively. The only interactions where P < 0.05 for at least one trait were RH x PAR and Genotype x PAR (Table S1).

The fractional contribution of source water (unenriched xylem water; *f*_sw_) was slightly greater in *Zmasft*mutants (0.22 ± 0.03) than wildtype (0.16 ± 0.03). Alternatively, for both genotypes *f*_sw_ was greater under 1200 compared to 300 μmol quanta m^-2^ s^-1^(0.27 ± 0.1 and 0.1 ± 0.02, respectively) (Table 1; Fig. 2a) and RH increased *f*_sw_ under 300 but not 1200 μmol quanta m^-2^ s^-1^ (Table 1; Fig. 2a). Transpiration rate (*E*; mol H_2_O m^-2^ s^-1^) was greater in the*Zmasft* double mutants than in wildtype across treatments (P < 0.0001), and this difference was most apparent under high light intensity. At high light intensity, *E* was higher than at low light intensity, and RH only affected *E* at low light intensity (Table 1; Fig. 2b). Similarly, stomatal conductance (*g*_s_; mol H_2_O m^-2^ s^-1^) was 44 % greater in*Zmasft* double mutant, and this genotypic difference was most apparent at high light intensity (Table 1).

**Fig. 2.**
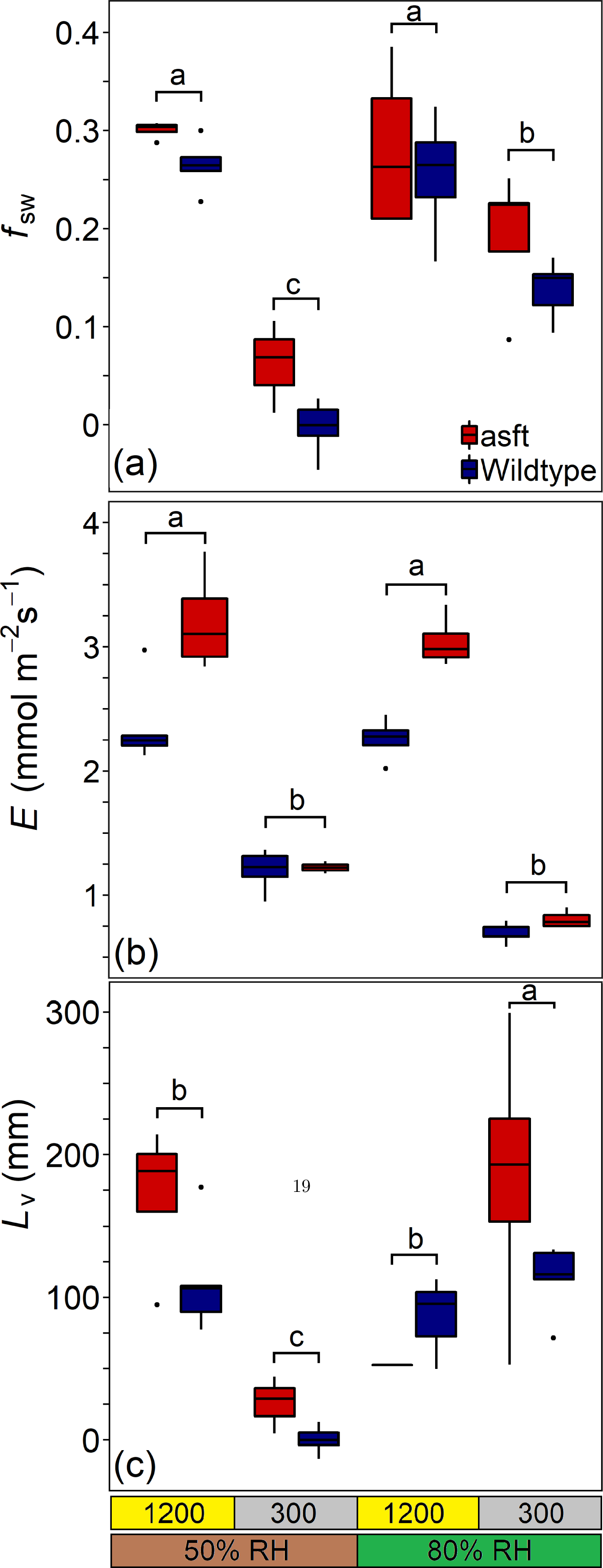

### Rice mixed linkage glucan mutants (cslf6-1 and cslf6-2) versus wildtype

Δ^18^O_e_ was greater in the wildtype (24.52 ± 1.00 *cslf6-1* (22.12 + −0.30 light (24.78 + −0.63 genotype (Table 2). Δ^18^O_LW_ was significantly higher in *cslf6-1* than wildtype and *cslf6-2* mutants, but higher Δ^18^O_LW_ in *cslf6-1* was not due to higher Δ^18^O_LW_ which was lower in *cslf6-1* than in wildtype and *cslf6-2* (Table 2; Fig. 3b). There was a higher proportion of enriched than unenriched water (lower *f*_sw_) in *cslf6-1* plants (0.36 ± 0.02) compared to wildtype (0.49 ± 0.02) and *cslf6-2* (0.48 ± 0.02) mutant (Table 2). In all genotypes, *f_sw_* was higher under high light than low light (0.48 ± 0.02 and 0.40 ± 0.02, respectively; Table 2; Fig. 4a). *E* and *g*_s_ were higher in wildtype (42 and 40 %, respectively) and *cslf6-2*(32 and 34 %, respectively) than in *cslf6-1* under high light, but there was no difference between genotypes under low light (Table 2). *E* and *g*_s_ significantly decreased under low light compared to high light in wildtype and *cslf6-2* but not *cslf6-1* (Table 2; Fig. 4b).

**Fig. 3.**
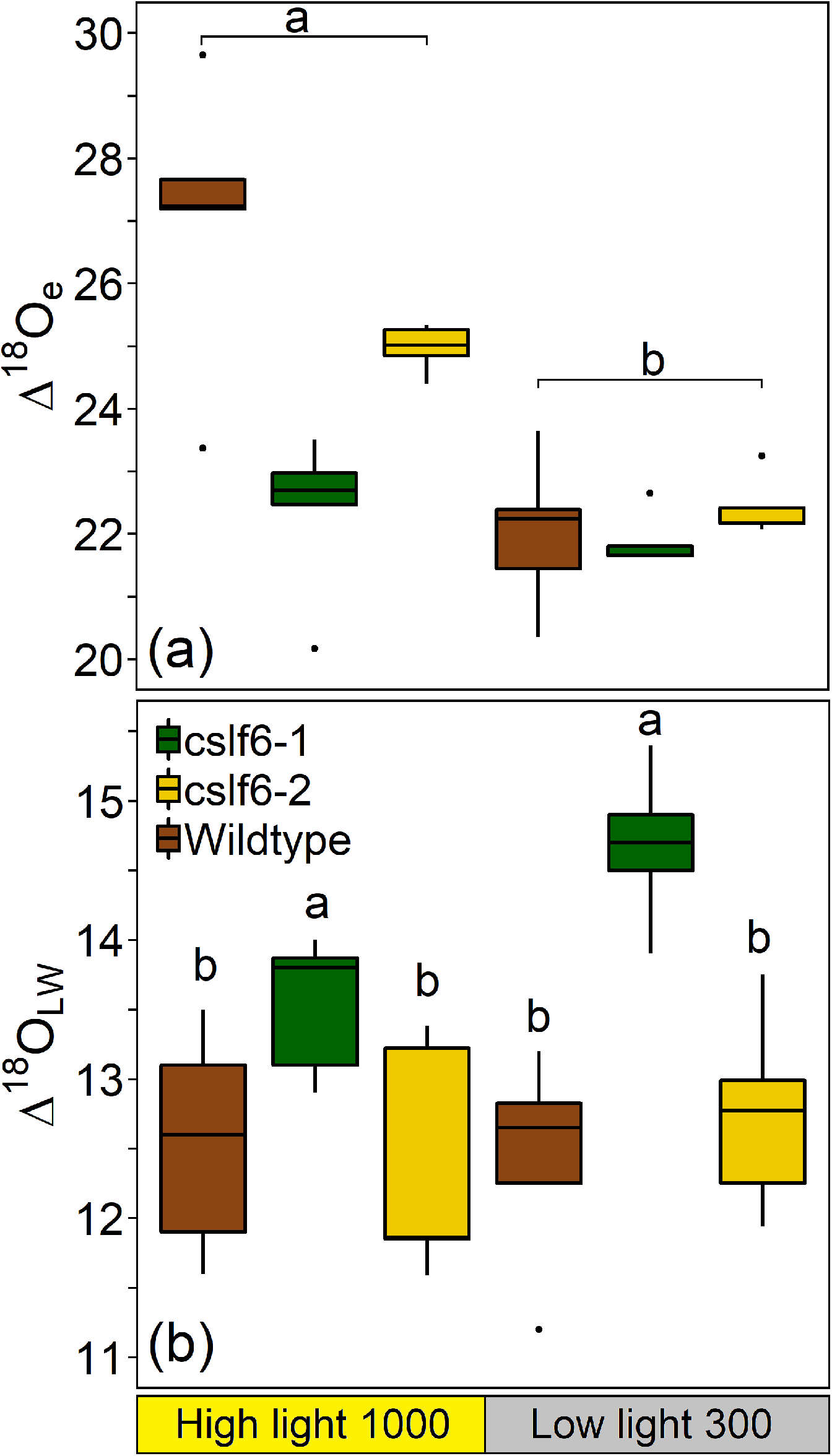

**Fig. 4.**
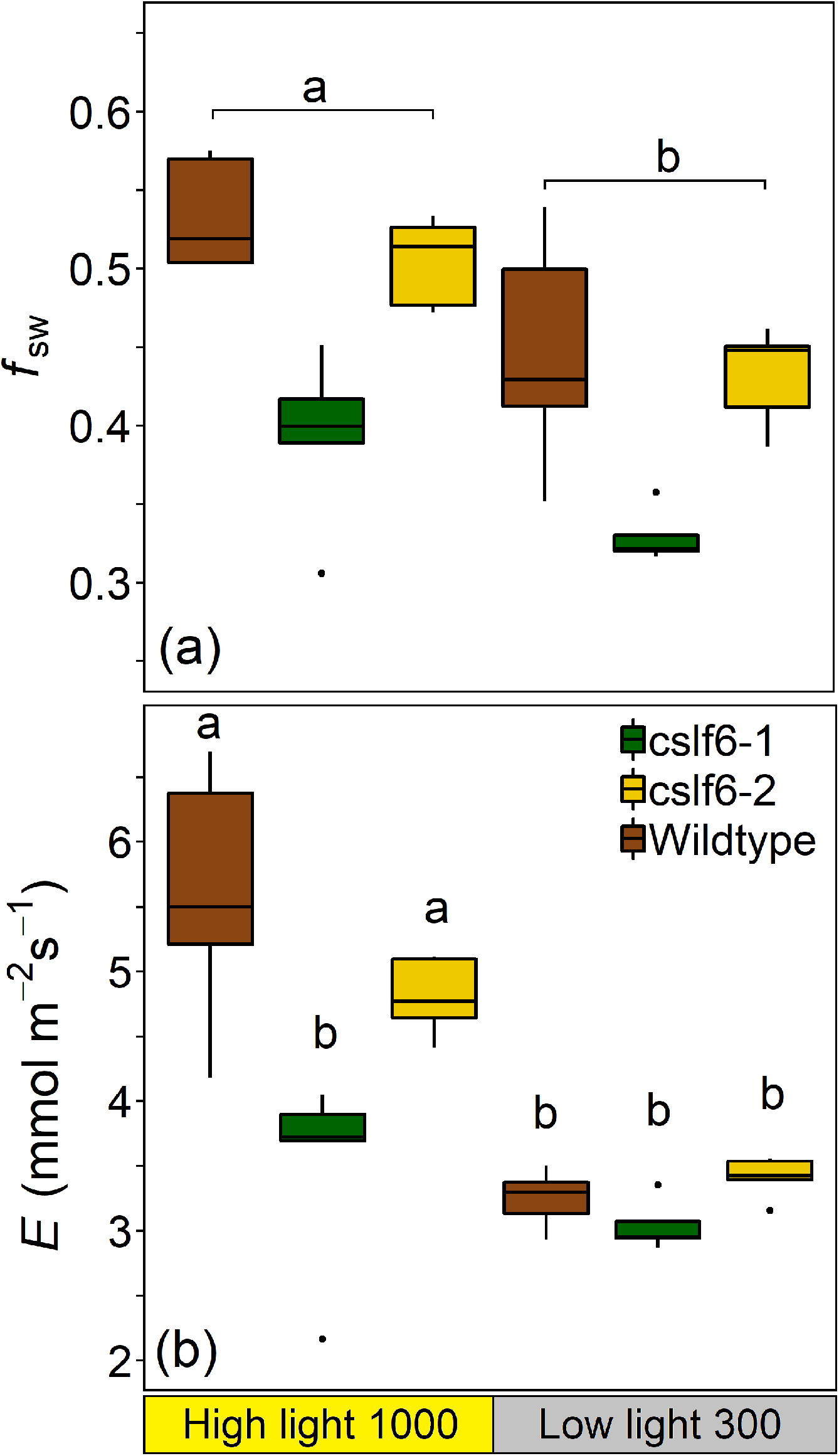

**Table 2.**
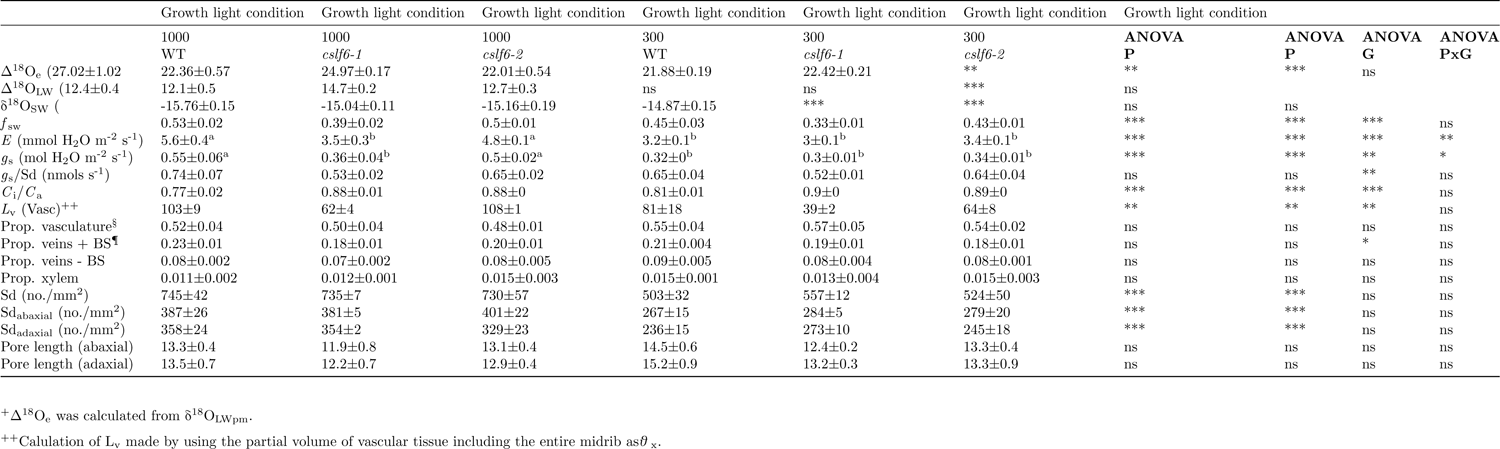
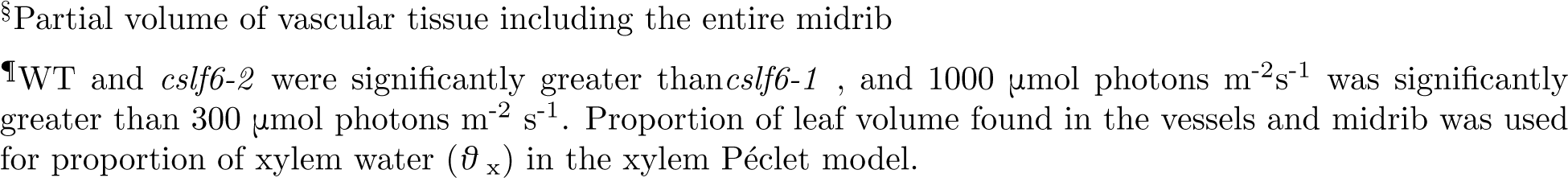
Leaf water isotope parameters for *O. sativa* wild type (Nipponbare), *cslf6-1* and *cslf6-2*, grown under 1000 and 300 photosynthetic active photon flux density (P, μmol photons m^-2^ s^-1^). *f*_sw_ is the proportion of unenriched water in bulk leaf water. Calculation of L_v_ made by using the partial volume of vascular tissue including the entire midrib as *θ*_x_. A two-way ANOVA was performed to test the significance: * *P* < 0.05, ** P < 0.01, *** P < 0.001. Tukey post hoc test represented when PxG interaction was significant. Values with the same superscripted letters are not significantly different. Growth light condition is represented by P (PAR) and genotype by G.

### Anatomical measurements in maize and rice

The partial volume of the leaf represented by vasculature (midrib and vascular bundles in the blade), veins (both including and excluding the bundle sheath cells), and xylem were not significantly different across genotypes in either maize or rice, except for the partial volume of veins including the bundle sheath in rice (Table 1 and 2). In rice, the partial volume of veins including the bundle sheath was slightly greater in WT (0.22 ± 0.02) than *cslf6-1* (0.18 ± 0.01) and *cslf6-2*(0.19 ± 0.01), but there was no difference in anatomy with growth light intensity (Table 2). The partial volume of total vasculature, veins (both including and excluding the bundle sheath), and xylem in maize (0.50 ± 0.01, 0.25 ± 0.01, 0.11 ± 0.01, 0.03 ± 0.01, respectively) and in rice (0.53 ± 0.04, 0.20 ± 0.03, 0.08 ± 0.01, 0.01 ± 0.02, respectively) showed that about half of the leaf was vasculature, and only a small fraction of the vasculature was conducting tissue (Table 2). In maize and rice, stomatal density (Sd) differed with growth light intensity where plants under high light had greater Sd (117.2 ± 1.5 and 737 ± 35, respectively) than those grown under low light (88.7 ± 1.4 and 528 ± 31, respectively) (Table 1 and 2). The Sd was slightly higher in leaves of *Zmasft* double mutants (105.7 ± 3.4) than wildtype (99.8 ± 3.9), but there were no differences across rice genotypes. Sd on both adaxial and abaxial sides of the leaf was significantly higher under high light, but neither was significantly different across genotypes (Table 1, S1). In maize, abaxial stomata subsidiary cells length was greater in wildtype grown under low light intensity (34.5 μm ± 2.0) than all other conditions (25.7 to 26.6 μm; Table 1, S1).

Changes in Δ^18^O_LW_ between plants grown in high and low light (HL-LL) co-varied with changes in *L*_v_, *E*, Sd, Sd_abaxial_, and partial volume of veins excluding the bundle sheath (P < 0.05) and partial volume of the total vasculature and *g*_s_ (P = 0.05; Table S2). The mean genotypic difference (HL-LL) in Sd and *E* formed significant linear relationships with Δ^18^O_LW_ (Fig. 5a, b), while *E* and Sd also formed a linear relationship (Fig. 5c). In contrast, the change in Δ^18^O_LW_ between plants grown at high and low light did not co-vary with changes in Δ^18^O_e_, mesophyll and bundle sheath cell wall thickness (bordering the IAS), leaf thickness, mesophyll cell layers, volume fraction of intercellular air spaces (IAS) per mesophyll area, mesophyll surface area exposed to IAS per unit leaf area (*S*_mes_), surface area of chloroplasts exposed to IAS per unit of leaf area (S_c_), and leaf dry mass per area [Table S2, these data were previously published in Ellsworth *et al*. (2018)].

**Fig. 5.**
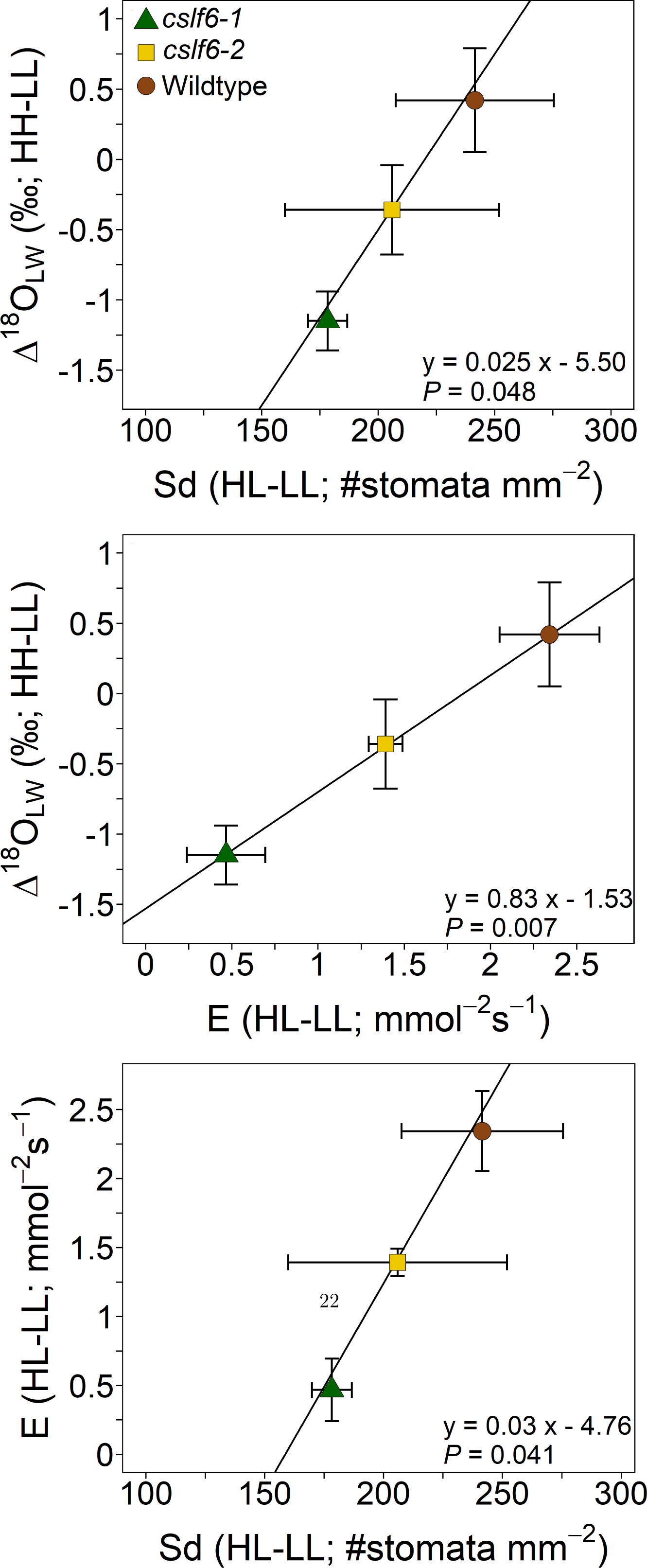

### Mesophyll and xylem P éeclet models in maize and rice

All genotypes of maize and rice formed significant positive relationships between *f*_sw_ and transpiration rate (*E*), suggesting a Péclet effect in these genotypes (Fig. 6a,b). When *f*_sw_ and *E* were fitted to the mesophyll and xylem Péclet model (Eqn. 10) to solve for the following three variables: mesophyll *L* (*L*_m_), xylem *L* (*L*_v_), and the proportion of xylem water (x), the model in maize converged, but the estimated value for *L*_m_ was essentially 0 (2 x 10^-17^). In rice the model failed to converge on a significant value for *L*_m_. Considering that *L*_m_ was estimated as ~0 in maize and did not converge on a significant value in rice, it was assumed that the mesophyll Peclet was nonexistent, and the location of the Peclet effect was in the xylem only (Eqn. 11) for maize and rice. In maize, the xylem Peclet model failed to converge on significant values of *L*_v_ and x when all data were used. However, when only data from 80 % RH were examined, *Zmasft* had a larger estimate of *L*_v_ (186 ± 86 mm) than wildtype (113 ± 47 mm; Table 3). At 50 % RH, the xylem Peclet model did not converge. In rice, *L*_v_ calculated from the xylem Péclet only model was smaller in *cslf6-1* than wildtype and *cslf6-2*. In each species, estimates of x were similar across genotypes, but x was twice as large in rice than maize (Table 3).

**Fig. 6.**
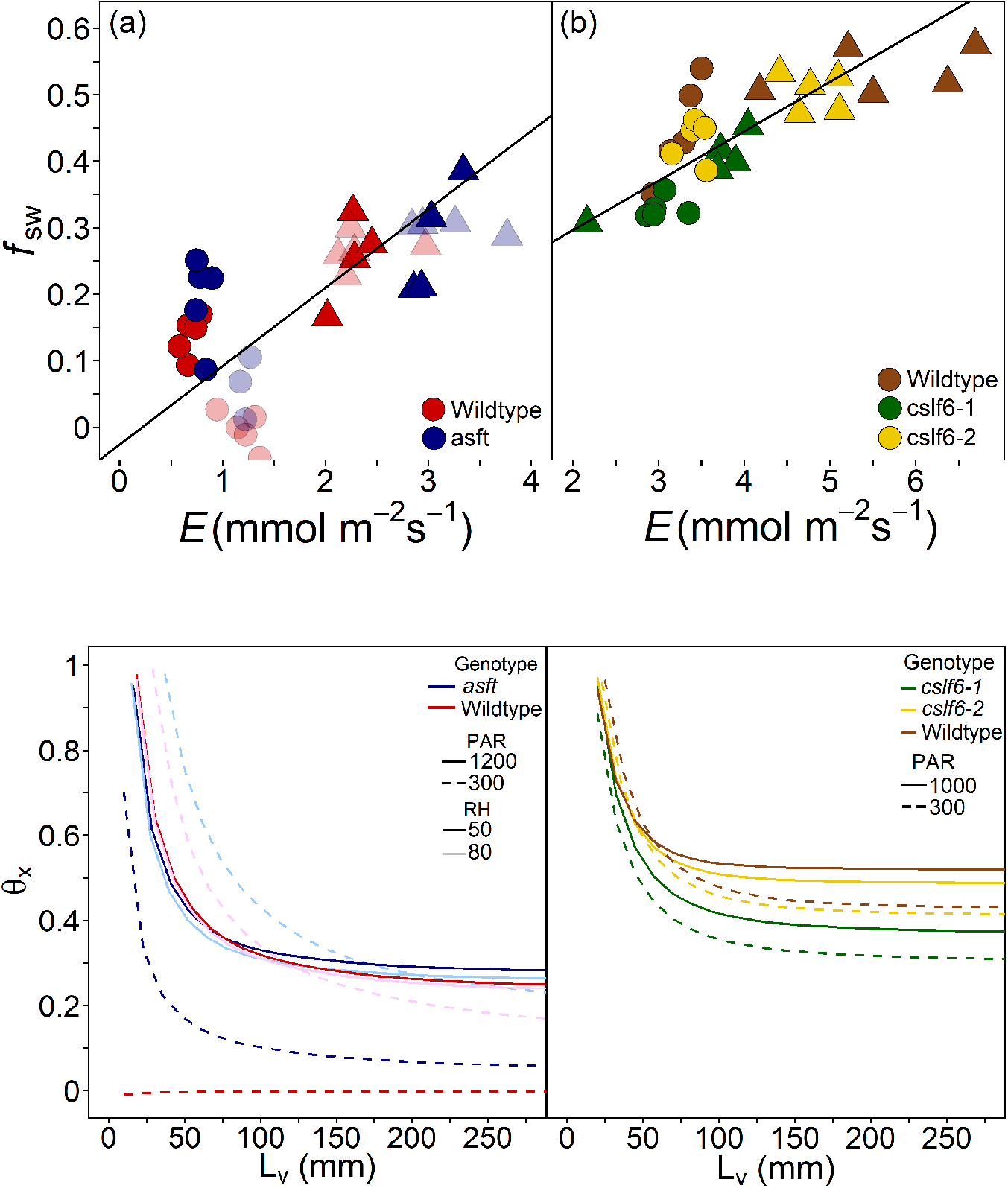

**Table 3.**
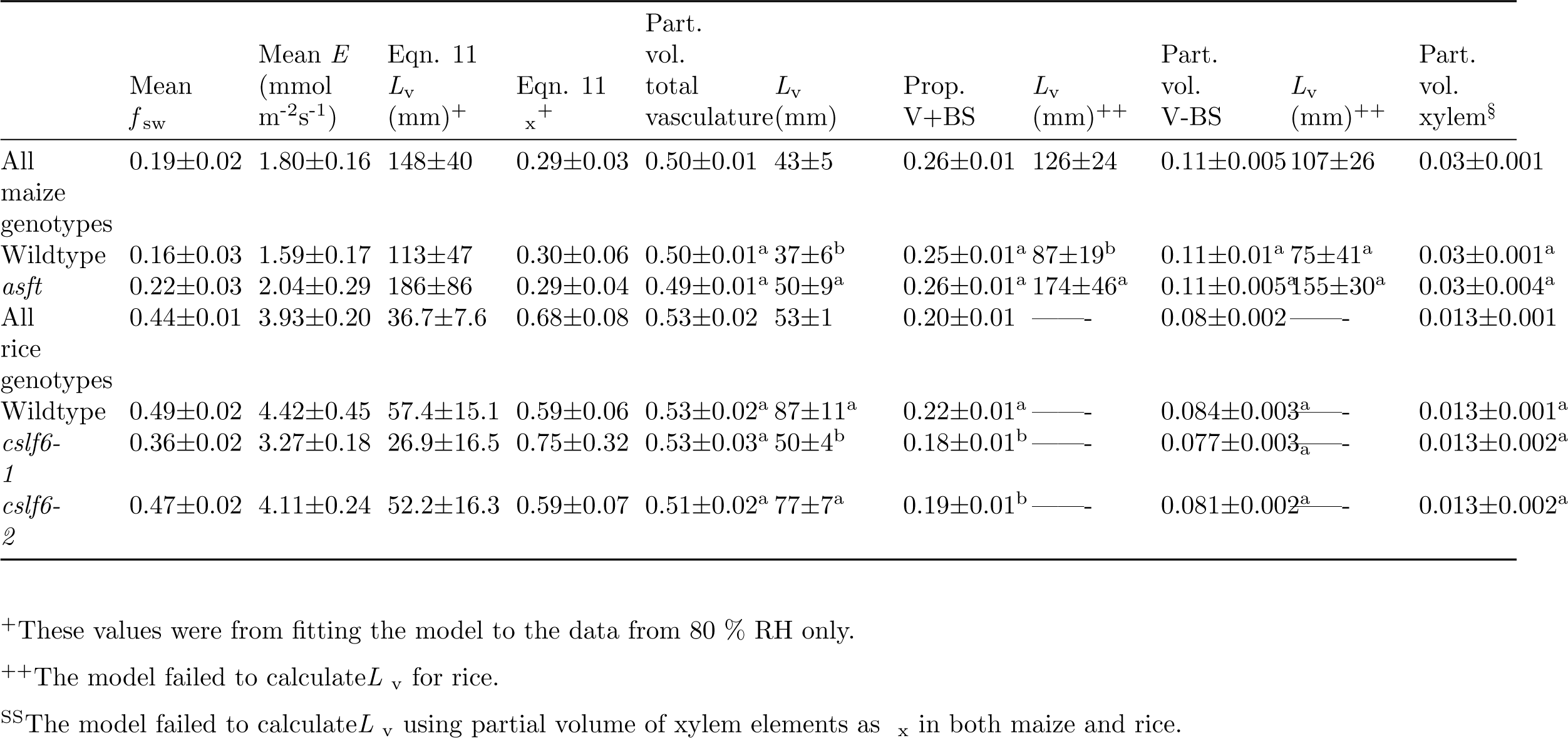
Xylem Péclet model (Eqn. 11) fitted to *f*_sw_ (1 - Δ^18^O_LW_/Δ^18^O_e_) and *E* data. Unknowns were *L*_v_ and x in the xylem Péclet model. Values are ± SE. *L*_v_ was calculated again when x was considered the partial volume of either total vasculature, veins including the bundle sheath (V + BS) or excluding the sheath (V - BS), or xylem. Means within each species and each column followed by different letters are significantly different.

*L*_v_ was estimated using the xylem Peclet model when the partial volumes of total vasculature, veins including and excluding BS were used as x in maize and only when the partial volume of total vasculature was used as x in rice. When the partial volume of leaf vasculature or vascular bundles was used as x in the xylem Péclet model, estimates of *L*_v_ were greater for *Zmasft* double mutants than WT, and this difference was most apparent at low light intensity (Table 1, 3). In rice, *L*_v_ was lower in *cslf6-1* than WT and *cslf6-2*. These results were similar to modelled *L*_v_ (solving for both *L*_v_ and x), except that *L*_v_ was considerably smaller when partial pressure of the total vasculature was used as x in maize. The partial volume of veins, including the bundle sheath, was most similar to *f*_sw_ (especially at high light intensity) and x values estimated from the xylem Péclet model in maize (Table 3). Rice, having larger *f*_sw_ values than maize, the partial volume of the total vasculature of rice was most similar to *f*_sw_ and x.

A sensitivity analysis of the interaction between *L*_v_ and x in the xylem Péclet model showed that x in rice was greater than in maize and that *L*_v_ had a large effect on x until it reached its minimum threshold in both maize and rice at approximately 75 mm (Fig. 7). Genotypic differences were only apparent at low light, where *L*_v_ and x were greater in *Zmasft* than in wildtype. In rice, *L*_v_ and x were lower in *cslf6-1* than wildtype and *cslf6-2* when *L*_v_ was greater than approximately 50 mm. The results between maize at 50 % RH and rice were similar in that x was greater at high light than low light across the range of *L*_v_ (Fig. 7).

## Discussion

Observations of Δ^18^O_LW_ values in maize and rice are consistent with the Péclet effect in the veins. Unlike the two-pool model where the *f*_sw_ is a fixed volume and not dependent on physiological factors (Roden & Ehleringer 1999; Roden *et al*. 2015; Song *et al*. 2015; Hirl *et al*. 2019), a positive relationship between *f*_sw_ and transpiration rate (*E*) was observed in all genotypes of maize and rice. According to the Farquhar and Gan model, Δ^18^O_LW_ is the accumulation of the Péclet effect in each tissue relative to the fraction of leaf water in the tissue, but several studies have found that the xylem water pool is the principal, if not the only, location of the Péclet effect (Helliker & Ehleringer 2000; Gan *et al*. 2002; Farquhar & Gan 2003; Gan *et al*. 2003; Holloway-Phillips *et al*. 2016). In support of this observation, we found that *L*_m_ estimates in the mesophyll and xylem Péclet model were essentially 0, indicating that the mesophyll Péclet effect was likely nonexistent. However, observed variation in Δ^18^O_LW_ across genotypes, species, and treatments was not related to the differences in the partial volume of the vasculature, meaning that variation in *f*_sw_ was influenced by anatomical and physiological factors regulating water movement and the relative importance of leaf water pools.

Anatomical differences between C_3_ and C_4_ grasses such as suberization of the bundle sheath regulate the movement of water out of the vasculature and through the mesophyll, contributing to variation in leaf water isotopic composition. For example, it is not clear which tissues should be included in the xylem water pool (x) of the xylem Péclet model, and x could include only the vein elements or include associated ground tissue such as the bundle sheath and the large quantity of parenchyma in the midrib (Gan*et al*. 2003). In C_3_ rice, the estimated x (and *f*_sw_) were relatively similar to the partial volume of the entire vasculature, which indicates that likely conducting and ground tissues associated with the vasculature together are isotopically one pool of water. Previous studies have made similar conclusions (Gan *et al*. 2003; Song *et al*. 2015; Holloway-Phillips *et al*. 2016). In contrast, the influence of xylem water in C_4_ maize may be more constrained than in rice in that x may be confined to the vascular bundles and may not necessarily include all ground tissue associated with the veins. This difference between C_3_ and C_4_ grass species may lie in that C_3_ plants have less suberized bundle sheaths, leading to greater outside xylem conductivity to support higher *E* and a greater isotopic influence of xylem water on the mesophyll water pool (Sonawane *et al*. in review; Kocacinar & Sage 2003). Therefore, the lower Δ^18^O_LW_ in C_3_ grass species than C_4_ species may depend on more than anatomical features such as interveinal distance (vein density) and include the facility of water movement out of the xylem and outside xylem conductivity.

### Suberin mutation and the Péclet effect

Both *E* and *L*_v_ were affected by the compromised ultrastructure of the bundle sheath cells and increased apoplastic permeability in *Zmasft* maize double mutants, especially at the interface of suberin lamellae and adjoining polysaccharide cell walls (Mertz *et al*. In review). Suberin is considered essential as a functional barrier to unrestricted diffusion of water across cell walls (Botha *et al*. 1982; Schonherr 1982; Mertz & Brutnell 2014). Consequently, more permissive water flow in the leaf of the *Zmasft* double mutants led to greater water flow out of the xylem as evidenced by greater *E* under high light and more water channels in and out of the xylem under low light, increasing the overall effective mixing length in the xylem (*L*_v_). The likely reason for mutation-related differences in *E* under high light and *L*_v_ under low light is an interaction between water demand and the lack of water channeling. Under high light *E* was 2-3.3x and 2.6-3.8x greater in wildtype and the double mutant relative to low light, respectively, so the principal water channels moved the vast majority of water out of the xylem, leading to similar measures of *L*_v_. In contrast, transpiration demand was low under low light, so greater apoplastic permeability allowed water to move through more tortuous channels, increasing the overall *L*_v_ in the *Zmasft* mutants. Therefore, channeling of water outside the xylem through suberin lamellae influenced the Péclet effect in leaf water by restricting water flow (lower *E*) and reducing water movement across the suberin lamellae (smaller *L*_v_). As has been seen previously, suberin and suberin lamellae play a role in the plant’s ability to regulate transpiration (Schreiber 2010; Vishwanath *et al*. 2015), but interestingly these differences are detectable with stable isotopes. Potentially, through the Péclet effect model Δ^18^O_LW_ can be used to detect and study anatomical differences and their physiological consequences on water movement in leaves.

### Mixed linkage glucan (MLG) and the Péclet effect

The genotypes varied in their physiological and isotopic response to growth light intensity. The genotypes (*cslf6-1*, *cslf6-2*), which are transposon insertion lines created in a Nipponbare background, did not always have the same phenotype (Upadhyaya *et al*. 2006; Vega-Sanchez *et al*. 2012). They were similar in most anatomical features and biochemical properties (Vega-Sánchez *et al*. 2012; Ellsworth *et al*. 2018). However, genotype *cslf6-1* had lower values than *cslf6-2* in several physiological traits such as *g_s_*, *E*, and both *A*_max_ and *J*_max_ on a per gram basis (Ellsworth *et al*. 2018) and *L*_v_ in this study. Because all plants were grown in the same physical space and under the same conditions such as temperature and relative humidity, variation in these traits across genotypes can be either attributed to pleiotropic effects on the cell wall or background effects as seen by the differences between Nipponbare and the isogenic WT siblings of these different lines (Vega-Sanchez *et al*. 2012).

The genotypic response of Δ^18^O_LW_ to growth light intensity can be explained within the Peclet effect theory by the corresponding response in stomatal density (Sd). Multiple studies have shown that Sd and stomatal size influence Δ^18^O_LW_ by modifying *E* and *L*, and that differences in Sd were sufficient to drive specieslevel differences in Δ^18^O_LW_ between mangrove and nearby freshwater species and *across Arabidopsis* stomatal density lines (Rosado *et al*. 2013; Sternberg & Manganiello 2014; Larcher *et al*. 2015; Liang *et al*. 2018). In this study, lower *L*_v_ in *cslf6-1* was likely due to an interaction between a genotype-driven reduction in *E* related to Sd and an increased mesophyll and xylem water exchange due to lower water flow, which reduced the mixing length between enriched mesophyll water and unenriched water in the xylem (Farquhar & Gan 2003; Gan *et al*. 2003). Therefore, changes in Sd regulated the relative differences in Δ^18^O_LW_, but Sd alone did not predict the magnitude of Δ^18^O_LW_(Rosado *et al*. 2013; Larcher *et al*. 2015). Conversely and similar to Sternberg and Manganiello (2014), higher Δ^18^O_e_ at high light in wildtype and *cslf6-2* did not result in higher Δ^18^O_LW_ because increased advective flow (*E*) and longer mixing length (*L*_v_) between xylem and mesophyll reduced back diffusion and the impact of Δ^18^O_e_ on Δ^18^O_LW_. The fact that the partial volume of the veins along with Sd, *L*_v_, and *E* co-varied with Δ^18^O_LW_ suggests that Δ^18^O_LW_ was significantly influenced by the entire water transport path through the leaf and, consequently, may trace anatomical, physical, and physiological factors affecting water movement.

### Conclusions

Cell wall and other anatomical properties can affect Δ^18^O_LW_ by manipulating *E* through altering water flow along the transpiration stream and *L*_v_ through altering water channeling out of the xylem and to the stomata. This influence is apparent in variation in Δ^18^O_LW_ between C_3_ and C_4_ grass species that can be attributed to differences in vasculature that limit water flow out of the vasculature in C_4_ species and may be an indication of differing hydraulic attributes. Studying the interactions of anatomical features and physiological traits in water movement through and outside of the xylem can shed more light on Δ^18^O_LW_ because either physiology or anatomy alone fails to capture the isotopic implications of water movement. Consequently, stable isotopes can be used to study this interaction. With the development of gene editing, Δ^18^O_LW_ can potentially be used to better understand gene-phenotype relationships of traits related to water movement in the leaf. For example, when studying genes associated with Sd, differences in water movement may not be discernable from measures of Sd *per se*, but nevertheless differences in Sd can influence Δ^18^O_LW_ and indicate a subtle response of Sd to genotype or environmental conditions. Further integration between leaf water transport through and outside the xylem with stable isotope theory will facilitate the development of a physiologically and anatomically explicit water transport model.

## Acknowledgements

Our research is supported by the Office of Biological and Environmental Research in the DOE Office of Science (DE-SC0008769) and by the Russian Science Foundation (16-16-00089) for N.K. (leaf anatomy study and anatomical traits analysis). We are grateful to the Core Facility Center “Cell and Molecular Technologies in Plant Science” of Komarov Botanical Institute and Franceschi Microscopy and Imaging Center of Washington State University for use of their facilities. Seed *Oryza sativa*, Nipponbare cultivar (wild type) and mutants*cslf6-1* and *cslf6-2* were kindly provided by Drs. Miguel E. Vega-Séanchez and Pamela C. Ronald, Joint BioEnergy Institute, Emeryville, California 94608 and UC Davis Genome Center and Plant Pathology, University of California, Davis, California 95616.

